# Individuals lack the capacity to accurately detect emotional piloerection

**DOI:** 10.1101/2023.07.19.549671

**Authors:** Jonathon McPhetres, Ailin Han, Halo H. Gao, Nicole Kemp, Bhakti Khati, Cathy X. Pu, Abbie Smith, Xinyu Shui

## Abstract

Piloerection (e.g., goosebumps) is an essential thermoregulatory and social signalling mechanism in non-human animals. While humans also experience piloerection—often being perceived as an indicator of profound emotional experiences—its comparatively less effective role in thermoregulation and communication might influence our capacity to monitor its occurrence. We present three studies (total N = 617) demonstrating participants’ general inability to detect their own piloerection events and their lack of awareness that piloerection occurs with a similar frequency on multiple anatomical locations. Participants over-reported piloerection events with only 31.8% coinciding with observable piloerection, a bias unrelated to piloerection intensity, anatomical location, heart rate variability, or interoceptive awareness. We also discovered a self-report bias for the forearm, contradicting the observation that piloerection occurs with equal frequency on multiple anatomical locations. Finally, there was low correspondence between self-reports of being “emotionally moved” and observed piloerection. These counterintuitive findings not only highlight a disconnect between an obvious physiological response and our capacity for self-monitoring, but they underscore a fascinating divergence between human and non-human species. While piloerection is vital in non-human organisms, the connection between piloerection and psychological experience in humans may be less significant than previously assumed, possibly due to its diminished evolutionary relevance.

**Statement of Impact:** This research reveals a striking dissociation between an obvious physiological response— piloerection—and human capacity for self-perception. While this highlights the limitations of relying on self-report measures, it also underscores an important divergence between humans and non-human species. We propose that the relation between piloerection and psychological experience in humans is less pronounced than in other species, potentially due to its diminished role in human evolution.

## Introduction

Piloerection—the erection of hair resulting from the contraction of the *arrector pili* muscle— is a highly conspicuous phenomenon observed across multiple animal species, including humans. In non-human species, piloerection serves as an important thermoregulatory and social-communicative mechanism (1). Avian species contract feather muscles for intricate mating displays (2), and mammalian species erect their body hair in response to perceived environmental threats (3). While humans lack sufficient body hair, rendering piloerection ineffective for the purposes of thermoregulation or social communication (4), humans do experience piloerection quite frequently.

Given the overt nature of piloerection and its utility in non-human organisms, one might expect humans to accurately monitor and report its occurrence. After all, piloerection is commonly perceived as an indicator of emotional experiences. Likewise, contemporary psychology predominantly presumes individuals possess accurate insight into their emotional experiences, often relying on self-reported data as the sole methodology (5). In contrast, because piloerection is assumed to be vestigial in humans, it may be that we lack the capacity to accurately monitor and report piloerection because it is void of any useful psychological or social information. Instead, humans have language to communicate internal states and we have clothes to assist with thermoregulation. Therefore, it is also reasonable to expect that humans would not monitor these physiological cues.

Nonetheless, the extent to which humans can accurately detect piloerection remains unexamined and, apparently, unquestioned. One reason for this is because much of the research on piloerection has relied on self-reports, rather than objective observation (1). In the studies reported here, we combine self-reporting with objective observation of piloerection on multiple anatomical sites, offering novel insight into individuals’ awareness and the nature of this physiological phenomenon. Understanding these dynamics is critical for refining our approach to interpreting data about human emotional and physiological experiences.

## Method

### Study 1

#### Participants

In a laboratory experiment, 90 subjects were recruited from the university participant pool and the surrounding community. There were 72 females and 18 males ranging in age from 18 to 50 (M = 21.08, SD = 5.33). All subjects reported being healthy, not smoking or drinking alcohol daily, not taking any medications, and were awake for at least two hours prior to the session. Participants were compensated with either course credit or cash payment.

#### Procedure

Participants arrived in a laboratory where they were connected to physiological recording equipment (BioPac MP160) and seated in a private cubicle at a computer running e-Prime (v2.0; Psychology Software Tools). Though not relevant to the results reported here, we also collected electrodermal activity, skin temperature, cardiac impedance, and blood pressure, and two saliva samples. Following a baseline period, they viewed 7 piloerection inducing videos in randomised order. Stimuli qualities and selection are not relevant to present report, but further details of the videos are available in the Supplementary Materials. Heart rate was monitored via Lead II electrocardiogram, piloerection was continuously recorded via high-definition video cameras (similar to (6)), and participants self-reported goosebumps via a button-press in real time.

The video recordings were analysed manually using BORIS (7) to accurately identify the beginning timepoint of each piloerection episode. Two coders manually coded each video, achieving 100% agreement; the first author then reviewed a subset of videos for quality checks. Using the MindWare EDA software (v. 3.2.9; MindWare Technologies LTD, Gahanna, OH), we searched for buttons presses within a 16-second symmetrical time-window around each piloerection event. Because piloerection can linger for long periods of time, a 16-second window was chosen so that we would overestimate (rather than underestimate) accuracy by allowing for participants to press the button at *the beginning or the end* of a given goosebumps episode.

### Study 2

#### Participants

Five hundred participants were recruited through Prolific in exchange for monetary compensation. There were 205 males and 291 females (4 identified as non-binary or preferred not to say) with a mean age of 40.4 (SD = 13).

#### Procedure

Participants accessed an online survey on Qualtrics where they were randomly assigned to watch one of three videos from Study 1. Videos were selected to be short to reduce participant time investment (see supplementary materials). In order to help reduce reports of “the chills” or other sensations, participants were told that we are interested in goosebumps, were given a description and images of goosebumps, and were asked to pay attention to their bodily experience while watching the videos.

After the video, participants were asked if they experienced goosebumps. Those who answered “Yes” were shown an image of a body and asked to click on the one area where they experienced the most intense goosebumps, followed by a second image for which they were asked to click on up to 10 other spots where they experienced goosebumps. Finally, they gave their age and sex and the survey was completed in about 5 minutes.

### Study 3

#### Participants

In Study 3, 30 participants were recruited from the psychology participant pool as well as from the surrounding community. There were 24 females and 5 males ranging in age from 18 to 50 (M = 19.93, SD = 0.88); demographics data for one participant was not recorded. All participants were healthy, they did not smoke or drink alcohol daily, were not taking any medications, and were awake for at least two hours prior to the session. Participants were compensated with either course credit or cash payment.

Video recording data for 3 participants was lost due to a computer error, precluding the quantification of piloerection events for those participants. Thus, these three participants are not included in the analysis presented here, reducing our total sample size to 27.

#### Method

Similar to Study 1, participants arrived in a laboratory where they were connected to physiological equipment and seated at a computer running e-Prime (v 3.0) in a private cubicle. All setup, equipment, and analysis procedures were the same as in Study 1. However, the experiment procedure differed slightly (see details in the supplement). Following the baseline period, participants were randomly assigned to watch one of 2 videos used in Study 1: “Mom” (n = 16) and “Singer” (n = 14) videos. During these videos, participants were instructed to press a button whenever they felt “emotionally touched or moved”.

## Results and Discussion

In Study 1, 57% (n = 52) experienced objective piloerection and 51% (n = 46) self-reported *goosebumps*. This represents the most effective set of piloerection-inducing stimuli reported in the physiological literature to date (1). However, only 31.8% of objective piloerection events mapped onto self-reported piloerection. Specifically, subjects over-reported experiencing goosebumps, and very few of those self-reports occurred at the same time as observed piloerection.

Accuracy was largely homogeneous: only 15 participants (16%) were accurate at a level greater than chance. A Welch’s t-test reveals that, compared to those with lower accuracy levels, those participants did not experience more piloerection events (*p* = .800, *d* = .10 [95% CI: -.50, .70]), nor did they press the button more times (*p* = .600, *d* = .23 [95% CI: -.39, .85]).

We first explored the discrepancy between self-reported and observed piloerection events. In total, there were 1,447 self-reported goosebumps and 1,012 objective piloerection events—an average of 31.4 (*SD* = 51.6) self-reports and 19.4 (*SD* = 28.6) objective piloerection events per person. This contradicts the hypothesis that participants were not engaged or were not following instructions. To test the possibility that piloerection episodes were undetected due to limited camera placement, we increased the number of monitored camera locations from one (n = 50) to four (n = 40). However, the difference in accuracy between single-camera (26.6%) and multi-camera (33.7%) settings was negligible, accounting for only 2% of variance in accuracy (pseudo-*R*^2^ = 0.02, *p* = 0.230): a very small effect size.

In Study 2 (N = 500), we examined whether participants’ biased attention towards specific body regions influenced self-reporting. After watching one of three piloerection-inducing videos in an online survey, participants clicked on an image of a human body to indicate where they experienced goosebumps. While we are unable to verify that participants truly experienced piloerection, these self-reports serve as a proxy for the locations that participants monitor. As seen in Figure 1, results revealed a strong bias for the forearm, which accounted for 41% of unique reports (k = 251). Put differently: the majority of people indicated the forearm as the single most intense location and, when given the opportunity to select any number of other locations, they simply selected the other forearm. However, laboratory data from Study 1 revealed that piloerection occurs with relatively equal frequencies across all anatomical sites (χ^2^ = 0.40, *p* = 0.900), further contradicting the common belief that piloerection is more common on the forearm (see Figure 1; also see supplement for additional analyses). Additionally, accuracy rate was similar across all anatomical locations (pseudo-*R*^2^ < 0.01): 30.6% on the dominant upper dorsolateral arm, 35.76% on the dominant dorsal calf, 32.9% on the left anterolateral thigh, and 30.65% on the right anterolateral thigh.

**Figure 1.**
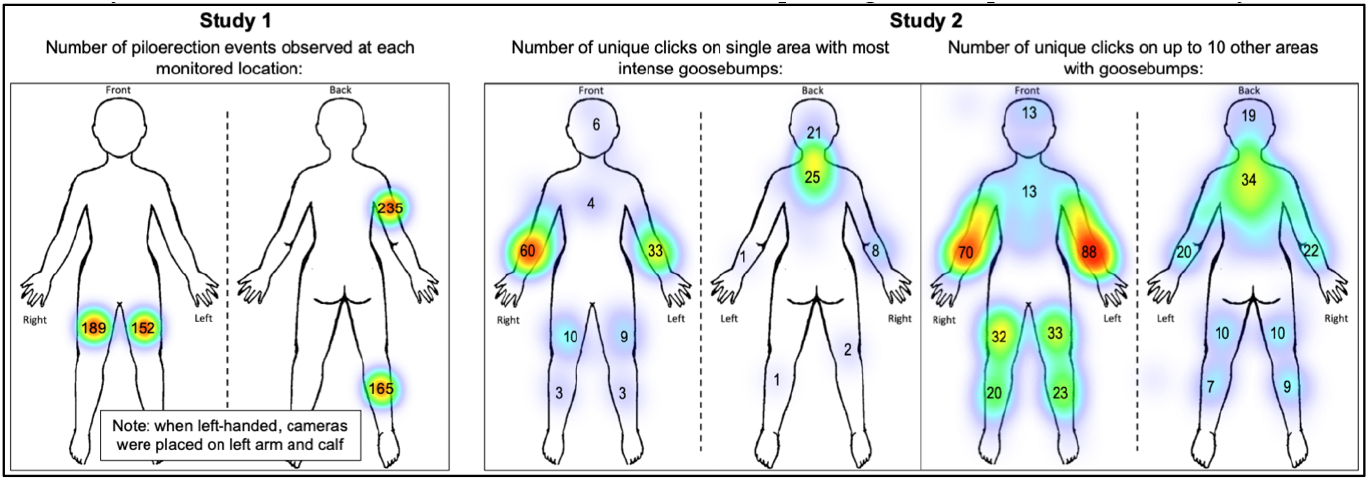
Whereas piloerection was observed with similar frequency on multiple anatomical locations in the laboratory, the forearm was the most common location of self-reported goosebumps in an online survey. *Note:* For Study 1, only counts for multi-camera participants are shown; For Study 2, N = 187 self-reported experiencing goosebumps at an average of 2.69 (mode = 1) locations; k = 2 erroneous clicks were excluded.

We then revisited data from Study 1, focusing instead on identifying those times when people were accurate. Contrary to the hypothesis that more intense episodes would be more noticeable, accuracy for the most intense episodes of piloerection remained below chance levels (40.1% on average; semipartial-*r*^2^ = 0.011, *p* < 0.001). Further, past research suggests that heart-rate variability moderates awareness of bodily sensations due to the correspondence between changes in the cardiopulmonary system and emotionally significant events (8). Emulating the approach taken in Lischke et al (8), we used two common measures of heart-rate variability—the proportion of successive heartbeat intervals exceeding 50ms (pNN50), and the root mean-square of successive differences between normal heartbeats (RMSSD). We also analysed each variability in two ways using separate mixed-effects regression models. First, we predicted accuracy levels with individual differences in baseline (i.e., resting) heart-rate variability. Second, we predicted variability measures (in 60-second epochs) around each piloerection episode with accuracy and baseline variability (e.g., a residual change model using mixed-effects regression). However, no correlations were found between accuracy and two heart-rate variability measures (all semipartial-*r*^2^ < .031, all *p* > .06).

Next, we tested the hypothesis that awareness of piloerection is an individual difference. We computed multiple sub-scales from the Multidimensional Assessment for Interoceptive Awareness (9) to approximate awareness of physiological sensations. Additionally, we added a single item “If I get goosebumps, I am aware of it.” However, a series of mixed-effects regression analyses indicate that no subscales, nor the single goosebumps item, were predictors of accuracy (all semipartial-*r*^2^ < .04, all *p* > .150). Additionally, those participants with accuracy levels greater than chance were no higher on any individual subscale (all *p*s > .200). In fact, 27 participants indicated that they would “Always” be aware of goosebumps (a score of 4 or 5 out of 5), although they were no more likely to be accurate than any other participant (Odds_log_ = .92, *p* = .330).

Finally, in Study 3 (N = 27) we considered the hypothesis that participants were conflating piloerection with “the chills” or other similar emotional experiences. Being “emotionally moved” is one of the experiences associated with the “chills” but which does not necessarily involve piloerection (10). In this study, participants used the button to self-report when they felt “emotionally moved or touched” while watching one of two piloerection-inducing videos. In total, 63% (N = 17) experienced piloerection—an average of 24 piloerection events per person—and 81% (N = 22) reported being emotionally moved or touched. However, only 24.6% of self-reports corresponded to piloerection. This again suggests a general difficulty in monitoring and reporting physiological experiences rather than conflation with similar physiological or emotional experiences.

## Conclusions

Overall, findings from these studies demonstrate a general inability to detect piloerection, as well as a lack of awareness regarding its occurrence across multiple body locations. These findings may be counterintuitive, as piloerection is frequently viewed as a hallmark of profound emotional experiences. However, the vestigial nature of piloerection in humans suggests that it should hold limited significance in the context of individual or social experiences. This may be one reason why it is not accurately monitored or reported.

This dissociation is important for understanding human emotional responses. A recent review (1) also discussed the lack of clear correlations between piloerection and self-reported emotions (see also (11). Therefore, either piloerection is not as closely tied to emotions as previously believed, or humans may not be as adept at identifying their emotional responses as assumed. Similarly, the apparent bias towards reporting piloerection on the forearm may shed light on human’s bodily awareness. The upper body is used for communicative purposes (12), whereas the legs are not usually a focal point in social situations. It may be a better use of cognitive resources to direct attention towards the arms, thereby leading to a bias where piloerection is ignored on other parts of the body.

A final point to address pertains to the subjective experience reported by participants—i.e., the self-reported goosebumps. There were a significant number of self-reports, suggesting that participants were experiencing some form of physiological or psychological event. It is conceivable that participants lack the perceptual acuity to distinguish between various physiological sensations, though a more plausible hypothesis is that this indicates the presence of a selection bias (e.g., survivorship bias). Given the emotional nature of the stimuli, participants likely experienced an emotional response. At times, emotions may seem to coincide with sensations that lack a physiological manifestation—for example, “the chills” or “tingles down the spine.” Operating under the common assumption that specific emotions correlate with piloerection, participants may infer that piloerection has taken place and report it. On occasion, this might coincide with an actual piloerection event, serving to reinforce the presumption that piloerection indicates specific emotions but ignoring the unnoticed piloerection events. Interestingly, scientists are familiar with, but not immune to, such biases. This is illustrated by nearly two decades of literature referring to piloerection as an indicator of awe despite the absence of any empirical evidence (11).

In conclusion, these findings highlight one important divergence between humans and non-human animals. There are many examples of physiological and psychological attributes shared between humans and non-human animals, including the fundamental structure of the nervous system which allows us to respond to changes in the external environment. It is easy to focus only on similarities between our species. In this case, however, piloerection serves as an example of how evolutionary pressures have contributed to the unique development of humans, leading us down a divergent evolutionary trajectory and distinguishing us from our non-human relatives.

## Ethics

Ethical approval was obtained from the Psychology Department Ethics Board at Durham University: PSYCH-2021-12-06T16_03_28-mqbg73

## Data Availability

Data are available on the open science framework at the following link: https://osf.io/ps2fq/?view_only=0c2ab9cd5beb410c9f48419351a1b8a1

## Notes

The human body outline depicted in Figure 1 was created by Thuy-vy Nguyen, who holds the copyright to the image. It is reproduced here with permission.

